# Validation and delineation of a locus conferring Fusarium crown rot resistance on 1HL in barley by analysing transcriptomes from multiple pairs of near isogenic lines

**DOI:** 10.1101/628420

**Authors:** Shang Gao, Zhi Zheng, Jonathan Powell, Ahsan Habib, Jiri Stiller, Meixue Zhou, Chunji Liu

## Abstract

**Background:** *Fusarium* crown rot (FCR) is a chronic and severe disease in cereal production in semi-arid regions worldwide. One of the putative quantitative trait locus (QTL) designated as *Qcrs.cpi-1H* has been previously mapped on chromosome arm 1HL in barley.

**Results:** In this study, five pairs of near-isogenic lines (NILs) targeting the 1HL locus were developed. Analysing the NILs found that the resistant allele at *Qcrs.cpi-1H* significantly reduced FCR severity. Transcriptomic analysis was then conducted against three of the NIL pairs, which placed the *Qcrs.cpi-1H* locus in an interval spanning about 11 Mbp. A total of 56 expressed genes bearing SNPs were detected in this interval, which would facilitate detailed mapping as well as cloning gene(s) underlying the resistance locus. Also, five differentially expressed genes (DEGs) bearing non-synonymous SNPs were identified in the interval. Differences in DEGs regulated by *Qcrs.cpi-1H* those by *Qcrs.cpi-4H* (another known locus conferring FCR resistance) indicate that different mechanisms could be involved in their resistance.

**Conclusion:** NILs developed in this study and the transcriptomic sequences obtained from them did not only allow the validation of the resistance locus *Qcrs.cpi-1H* and the identification of candidate genes underlying its resistance, they also allowed the delineation of the resistance locus and the development of SNPs markers which formed a solid base for detailed mapping as well as cloning gene(s) underlying the locus.

## Background

Fusarium crown rot (FCR), caused mainly by *F. pseudograminearum*, is a severe and chronic disease of cereals in semi-arid cropping regions worldwide [1, 2]. To reduce FCR damage, several agronomic measures have been developed. They include crop rotation and stubble management [3, 4]. These practices can reduce the impact of FCR in certain circumstances but are not always useful due to economic and practical requirements [5]. It has long been recognised that growing resistant varieties is an essential component to effectively manage this disease [6].

Similar to those for other diseases, identifying QTL conferring resistance and transferring them into elite genotypes are also used in breeding for FCR-resistant varieties in wheat and barley [7, 8]. Up to date, four QTL conferring FCR resistance have been reported in barley [9]. They locate on chromosome arms 1HL [10], 3HL [11], 4HL [12] and 6HL [13], respectively. Similar to those noticed in wheat [14, 15], strong interactions between FCR severity and other characteristics including flowering time [12, 16] and plant height [11, 17] have also been detected in barley. The FCR resistance locus on chromosome arm 3HL in barley also co-locates with gene(s) controlling spike structure [18]. Results from previous studies also showed that water availability affects FCR development [19].

The interactions between FCR severity and other characteristics indicate that QTL detected through mapping can only be treated as putative. The effectiveness of a QTL detected from segregating populations needs to be validated. Near isogenic lines (NILs) have been used widely in validating QTL for various characteristics [20, 21]. They were also used to validate QTL conferring resistance to FCR in cereals [22, 23].

Different from the main focus of detecting differentially expressed genes (DEGs) when the technique was initially introduced [24, 25], transcriptomic analysis is now also widely used to uncover genetic markers for various purposes [26, 27]. Combined with the use of NILs, distributions of variations detected from transcriptomic sequences have been exploited effectively in validating QTL and obtaining markers for fine mapping targeted loci [28-30].

In the study reported here, NILs were developed and used to validate the QTL conferring FCR resistance on 1HL. Transcriptomic sequences were then obtained from three pairs of the NILs. Transcriptomic responses mediated by the 1HL locus were analysed and the results were compared with those identified for another FCR resistance locus on 4HL [12, 29]. Shared SNPs detected from the transcriptomic sequences among the NIL pairs were used to further delineate the QTL interval and identify candidate genes underlying the resistance locus on 1HL.

## Materials and methods

### Development of near isogenic lines

The heterogeneous inbred family (HIF) method [31], combined with the fast-generation technique [32], was used to develop NILs targeting the 1HL locus (*Qcrs.cpi-1H*). Plants were raised in glasshouses at Queensland Bioscience Precinct (QBP) in Brisbane, Australia. Heterozygous plants were identified from two segregating populations, ‘Locker//AWCS079/AWCS276’ and ‘Commander//AWCS079/AWCS276’, using the SSR marker *WMC1E8.* This marker was one of those linked closely with *Qcrs.cpi-1H* identified from QTL mapping [10]. Primer sequences of the marker were: forward 5’-TCATTCGTTGCAGATACACCAC-3’; and reverse 5’-TCAATGCCCTTGTTTCTGACCT-3’. The identified plants were self-pollinated for eight generations and a single pair of putative NILs was then selected from each of the original heterozygous plants.

### FCR inoculation and assessment

FCR inoculation was conducted in the controlled environment facilities (CEFs) at Queensland Bioscience Precinct, Brisbane. Four inoculation trials, each with two replicates containing fourteen seedlings per isolines, were conducted against the putative NILs using a highly aggressive isolate of *Fusarium pseudograminearum* (*Fp*: CS3096). This isolate was collected in northern New South Wales and maintained in the CSIRO collection [33]. Procedures used for inoculum preparation, inoculation and FCR assessment were based on those described by Li et al. [34]. Briefly, seeds were surface-sterilized by treating with 2% hypochlorite solution for 10 min and then thoroughly rinsed with distilled water for four times. The seeds were then germinated on three layers of filter paper saturated with water in petridishes. Newly germinated seedlings (with coleoptile lengths ranging from 0.5 to 1.0 cm) were inoculated by immersing in *Fusarium* spore suspension (or water for controls) for 1 min. Two treated seedlings were sown in a 4cm × 4cm square punnet (Rite Grow Kwit Pots, Garden City Plastics, Australia) containing autoclaved potting mix. Fifty-six punnets were placed in a plastic seedling tray for easy handling. Inoculated seedlings were kept in CEFs. Settings for the CEFs were: 25/16(± 1) °C day/night temperature and 65%/85% day/night relative humidity, and a 14-h photoperiod with 500 mol m-2 s-1 photon flux density at the level of the plant canopy. Plants were watered only when wilt symptoms appeared. FCR severity for each plant was assessed with a 0-5 scale, where “0” standing for no symptom and “5” representing whole plant necrotic [34]. Disease indices (DI) was calculated for each line following the formula of DI = (∑_n_X / 5N) × 100, of which, *X* is the scale value of each plant, *n* is the number of plants in the category, and *N* is the total number of plants assessed for each line. The difference between the isolines possessing the resistant and susceptible allele for each of the putative NIL pairs was assessed with the student *t* test.

### RNA extraction and sequencing

Samples for RNA sequencing were obtained from three pairs of the NILs. Inoculation was conducted with either the F. pseudograminearum isolate (Fp-inoculation) or distilled water (mock) following the protocol described above. Three biological replications were conducted for every isolines. Each replication consists of seven seedlings. Tissues for RNA extraction were collected by cutting the shoot bases (2 cm) at 4 days post inoculation (dpi) and snap-frozen in liquid nitrogen and kept at – 80 °C until processed. The time point for sampling was selected based on a previous study [29].

A total of 36 samples were obtained from the six isolines. Samples were crushed into fine powder and RNA extraction was conducted using an RNeasy plant mini kit (Qiagen, Hilden, Germany) according to manufacturer’s instructions (including DNase-I digestion). The yield and purity of RNA samples were measured using a Nanodrop-1000 Spectrophotometer. The integrity of all RNA samples was assessed by running the total RNA on 1% agarose gels. RNA sequencing was carried out by the Australian Genome Research Facility Ltd (Parkville, Victoria, Australia) and 100-bp paired-end reads were produced using the Illumina Hiseq-2000. Four technical replicates were run for each of the 36 RNA-seq libraries.

### Transcriptomic analyses

Commands used for trimming raw data and analysing trimmed reads were described by Habib et al. [29]. FastQC (version 0.11.2) was used as a preliminary check for PHRED scores. Raw reads were trimmed using the SolexaQA package (version 3.1.3) with a minimum PHRED quality value of 30 and minimum length of 70 bp. TopHat2 (version 2.0.13) [35] was used to map filtered reads to the ‘Morex’ genome which is now widely used widely as the reference for barley [36].

### Differential gene expression analysis

Cufflinks (version 2.0.2) [35] was used to assemble the mapped reads. Differentially expressed genes (DEGs) were identified with Cuffdiff from the Cufflinks tool package with high-confidence genes annotated in the ‘Morex’ genome. Fragments per kilobase of exon per million mapped reads (FPKM) was applied for each transcript to represent the normalized expression value. The fold change in gene expression was calculated according to the equation: Fold Change = log_2_ (*FPKM*_*A*_/ *FPKM*_*B*_).

Pairwise comparisons were conducted between different treatments for the same isoline (S^M^_v_S^I^ and R^M^_v_R^I^) and between isolines under *Fp-*inoculation (S^I^_v_R^I^) or mock-inoculation (S^M^_v_R^M^). ‘M’ stands for ‘mock-inoculation’, ‘I’ for *Fp*-inoculation, ‘S’ for susceptible isolines, and ‘R’ resistant isolines. DEGs were determined with the adjusted *p*-value threshold of ≤ 0.05 and log_2_ fold change of ≥ 1 or ≤ −1 or ‘inf’ (where the FPKM value in one dataset is zero and the other is not). ShinyCircos was used to visualize DEGs on genomic level [37]. Venny 2.0 was used for Venn diagram analysis [38].

### Validation of differentially expressed genes using qRT-PCR

Three genes (*HORVU1Hr1G092240, HORVU1Hr1G092250* and *HORVU1Hr1G092300*; primers listed in Table S5) were selected from the identified DEGs for validation. Quantitative real-time PCR (qRT-PCR) was used for validation with the actin protein gene as the internal housekeeping reference (forward primer: 5’-GCCGTGCTTTCCCTCTATG-3’; reverse primer 5′-GCTTCTCCTTGATGTCCCTTA-3′). Inoculation, tissue sampling and RNA extraction were carried out using the aforementioned methods. Three biological replicates, each with two technical replications, were used for each genotype-treatment sample per isoline.

The procedures for synthesising cDNA and qRT-PCR were conducted following the methods described by Ma et al. (2013). The relative fold changes were calculated using the comparative CT method (2^-ΔΔCT^). The average value of the two technical replications was used to represent the biological replicate for each of the samples.

### SNP calling and nonsynonymous variation identification

For each genotype, all six sequence files (three biological replicates by two treatments) were concatenated after removing low-quality sequences. The concatenated files were then aligned to the ‘Morex’ genome using Biokanga align [39] with a maximum of two mismatches per read. SNPs between the ‘R’ and ‘S’ isolines of each NIL pair were identified using the Biokanga snpmarkers [39] with a minimum 80% score (the percentage of a given nucleotide at an SNP position is at least 80% in the ‘R’ or ‘S’ isoline). The SNPs were annotated using snpEff 4.3q [40] and the variant database was built based on the Morex genome and its annotation file [36].

### Gene annotation and gene ontology (GO) term enrichment analysis

BLAST, mapping and annotation steps were performed using the standard parameters in BLAST2GO [41]. DEGs identified from all comparisons were separated into up-regulated and down-regulated ones and subjected to singular enrichment analysis using agriGO [42].

### Comparison of the DEGs detected in this study with those from another FCR resistance locus on 4HL

Transcriptomic data from NILs targeting the FCR locus on 4HL were obtained from an earlier study [29]. Methods used between these two studies, including inoculum preparation, seedling age used for inoculation, conditions used for plant growth and sampling time for RNA sequencing, are all the same. The transcriptomic data used for comparison with those obtained in this study, including 4H_NIL1, 4H_NIL2 and 4H_NIL3, were downloaded from NCBI BioProject ID: PRJNA392021.

## Results

### Development and validation of NILs targeting the FCR resistance locus on 1HL

Eight heterozygous plants were initially selected from the two segregating populations based on the profiles of the SSR marker *WMC1E8*. A single pair of putative NILs was obtained from each of the heterozygous plants. Significant difference in morphology between any pairs of the putative ‘R’ and ‘S’ isolines was not observed. Significant difference in FCR severity was detected between the isolines for five of the eight putative NIL pairs. As expected, the isolines carrying the resistant allele from the donor parent AWC079 always gave much lower FCR severity than their counterparts (Table 1). The average DI for the ‘R’ isolines was 27.1, whereas it was 68.4 for the ‘S’ isolines. Three of the five NIL pairs with the largest difference in FCR severity, namely 1H_NILs: 1H_NIL1, 1H_NIL2 and 1H_NIL3, were selected and used for RNA-seq analysis.

**Table 1.** Difference in disease index between the resistant and susceptible isolines for the five NIL pairs targeting the 1HL locus conferring FCR resistance

| NIL<sup>a</sup> | Genetic Background | DI Mean<sup>b</sup> | Difference (%)<sup>c</sup> | P value<sup>d</sup> |
| --- | --- | --- | --- | --- |
| 1H_NIL1_R | Lockyer//AWCS079/AWCS276 F8 | 24.9 | 66.1 | <0.01 |
| 1H_NIL1_S |  | 73.7 |  |  |
| 1H_NIL2_R | Lockyer//AWCS079/AWCS276 F8 | 24.6 | 63.4 | <0.01 |
| 1H_NIL2_S |  | 67.3 |  |  |
| 1H_NIL3_R | Commander//AWCS079/AWCS276 F8 | 26.4 | 58.0 | <0.01 |
| 1H_NIL3_S |  | 62.9 |  |  |
| 1H_NIL4_R | Lockyer//AWCS079/AWCS276 F8 | 27.9 | 57.4 | <0.01 |
| 1H_NIL4_S |  | 65.5 |  |  |
| 1H_NIL5_R | Commander//AWCS079/AWCS276 F8 | 31.7 | 56.4 | <0.01 |
| 1H_NIL5_S |  | 72.7 |  |  |

<sup>a</sup>'R' represent isolines with the allele from the resistant parent 'AWC079' and 'S' isolines with an alternative allele from the susceptible parents.
<sup>b</sup>The mean of disease indices (DI value) observed from four trials for each isolate.
<sup>c</sup>Differences between DI values of 'R' and 'S' isolines.
<sup>d</sup>'P value' was generated with the student's *t* test.

### Transcriptome analyses

A total of 792 million quality reads were generate from the 36 samples (see the section of Materials and methods) with an average of 22 million reads per sample. The reads from each of the samples covered on average 21,571 high confidence (HC) genes (54.2% of all HC genes) based on the genome of Morex.

To analyse host response to *Fusarium* infection, differentially expressed genes (DEGs) were detected between *Fp*-and mock-inoculated samples of the same isoline. This analysis identified a total of 1,323 DEGs from the ‘R’ isolines and 2,083 from the ‘S’ isolines. The numbers of up-regulated genes were significantly higher than those down-regulated ones following *Fp*-inoculation (Table 2). Of the up-regulated genes, 144 were shared by all the three ‘R’ isolines and 370 by the three ‘S’ isolines (Fig. 1). Of the down-regulated genes, 17 were shared by the three ‘R’ lines and only 9 by the three ‘S’ lines. Expression patterns consistent with the RNA-seq analysis were obtained in the qRT-PCR analysis for each of the three genes assessed (Table S1).

**Table 2.** Number of DEGs identified from all pairwise comparisons

| NIL pair | Comparison<sup>a</sup> | Number of DEGs |  |
| --- | --- | --- | --- |
|  |  | Up-regulated | Down-regulated |
| 1H_NIL1 | R<sup>M</sup>_vs_R<sup>I</sup> | 226 | 60 |
|  | S<sup>M</sup>_vs_S<sup>I</sup> | 831 | 113 |
| 1H_NIL2 | R<sup>M</sup>_vs_R<sup>I</sup> | 962 | 132 |
|  | S<sup>M</sup>_vs_S<sup>I</sup> | 806 | 78 |
| 1H_NIL3 | R<sup>M</sup>_vs_R<sup>I</sup> | 910 | 117 |
|  | S<sup>M</sup>_vs_S<sup>I</sup> | 1585 | 252 |
| 1H_NIL1 | R<sup>I</sup>_vs_S<sup>I</sup> | 48 | 236 |
|  | R<sup>M</sup>_vs_S<sup>M</sup> | 225 | 123 |
| 1H_NIL2 | R<sup>I</sup>_vs_S<sup>I</sup> | 51 | 459 |
|  | R<sup>M</sup>_vs_S<sup>M</sup> | 178 | 89 |
| 1H_NIL3 | R<sup>I</sup>_vs_S<sup>I</sup> | 249 | 132 |
|  | R<sup>M</sup>_vs_S<sup>M</sup> | 80 | 71 |

<sup>a</sup>'M' stands for 'mock-inoculation', 'I' for *Fp*-inoculation, 'R' resistant isolines and 'S' for susceptible isolines

**Fig. 1.**
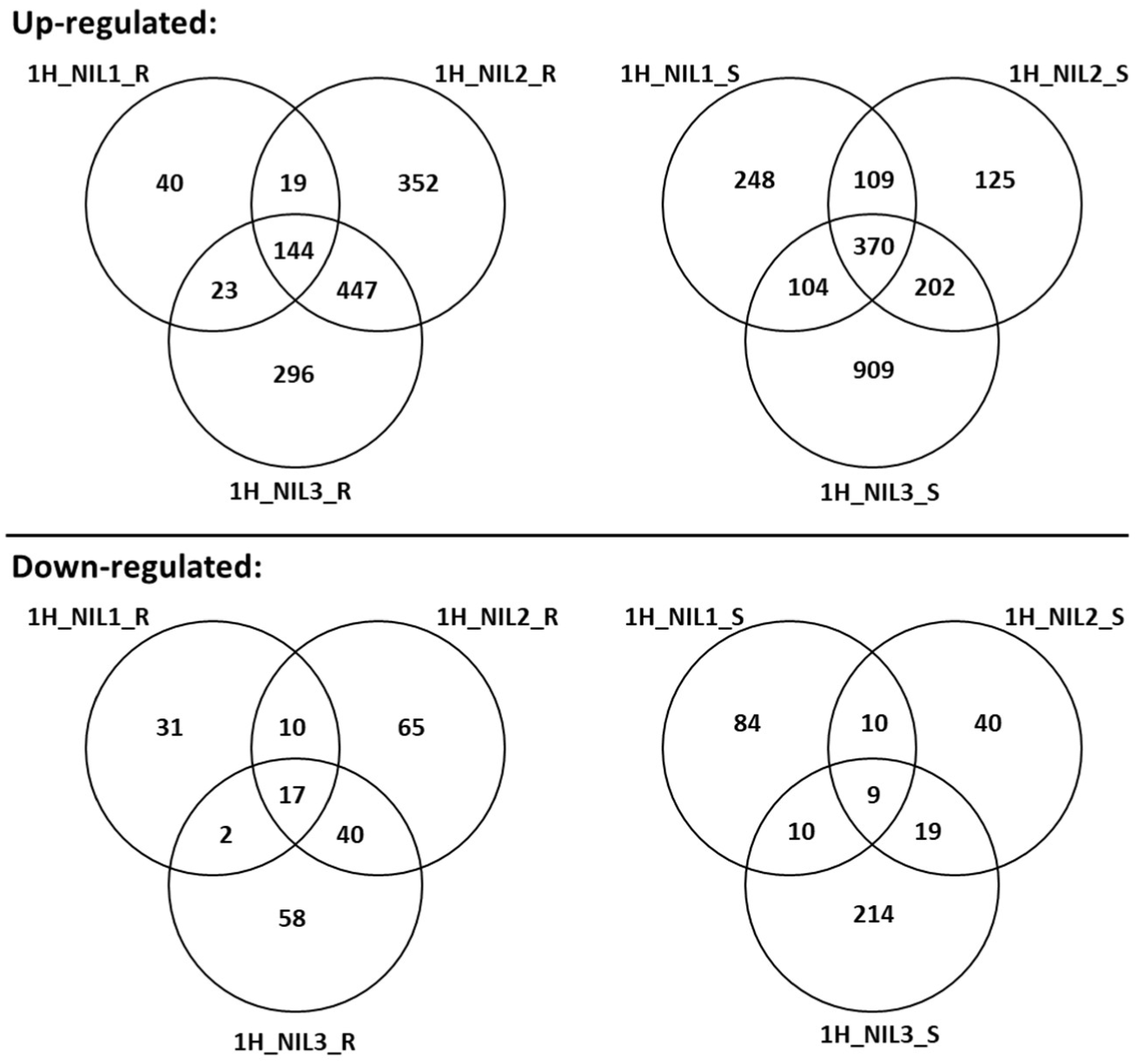
DEGs for each of the 1H_NIL pairs following *Fp*-and mock-inoculation (R^M^_vs_R^I^ and S^M^_vs_S^I^). Venn diagrams in upper panel show the numbers of up-regulated DEGs in each ‘R’ (left) and ‘S’ (right) isolines. Venn diagrams in lower panel show the numbers of down-regulated DEGs in each ‘R’ (left) and ‘S’ (right) isolines. DEGs were determined with the threshold of FDR ≤ 0.05 and |log_2_ fold-change|≥ 1 or ‘inf’ (one of the comparative objects did not express and the other did)

To assess transcriptomic responses to FCR infection mediated by *Qcrs.cpi-1H*, we compared DEGs between the ‘R’ and ‘S’ isolines. These comparisons found that a total of 303 genes were up-regulated and 790 down-regulated from the *Fp*-inoculation treatment (Table 2). Only 4 of the up-regulated genes and 2 of the down-regulated ones were shared by all three NIL pairs (Fig. 2). Of the DEGs identified from the mock-inoculated samples, 440 were up-regulated and 283 down-regulated (Table 2). Ten of the up-regulated and 3 down-regulated ones were shared across all the three comparisons (Fig. 2).

**Fig. 2.**
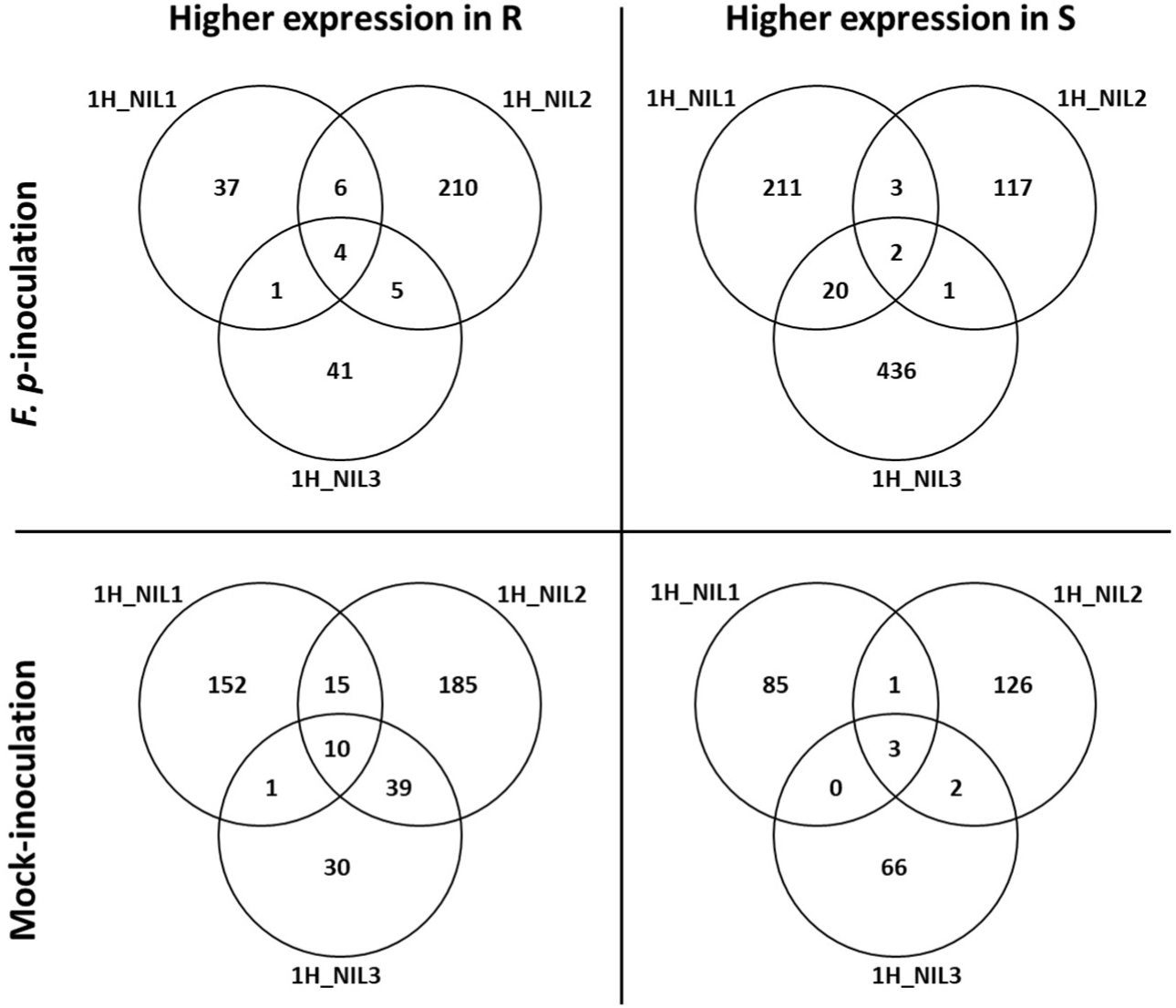
DEGs between ‘R’ and ‘S’ isolines under *Fp-* (R^I^_vs_S^I^) or mock-inoculation (R^M^_vs_S^M^). Venn diagrams show the numbers of DEGs which up-regulated in ‘R’ (left) or ‘S’ (right) isolines under *Fp-* (up) or mock-inoculation (down). DEGs were determined with the threshold of FDR ≤ 0.05 and |log_2_ fold-change|≥ 1 or ‘inf’ (one of the comparative objects did not expressed and the other did)

### SNPs between the ‘R’ and ‘S’ isolines across the three 1H_NIL pairs

In total, 2,753 non-redundant homozygous SNPs were detected between the ‘R’ and ‘S’ isolines. The number of SNPs detected from 1H_NIL2 was more than twice compared with those detected from either of the other two NIL pairs. Of these SNPs, 293 were common among the three pairs of the 1H_NILs. As expected, the majority of the SNPs shared among the three NIL pairs located at the distal end of chromosome arm 1HL where *Qcrs.cpi-1H* resides (Fig. 3). They spanned a physical distance of ∼ 11.0 Mbp (Fig. 4a).

**Fig. 3.**
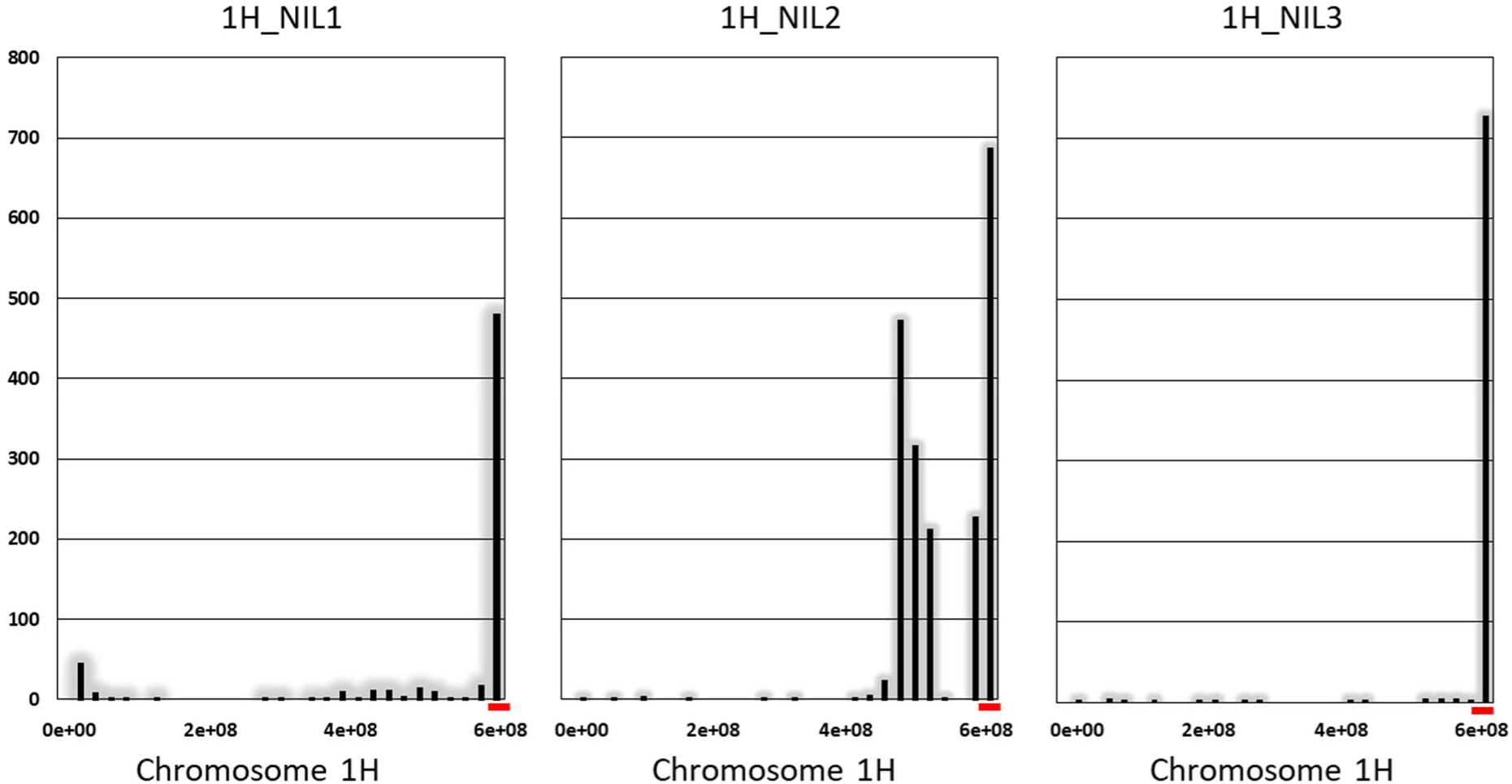
Distribution of SNPs in the expressed genes along chromosome 1H in three pairs of the 1H_NILs. Vertical axis shows number of SNPs. Horizontal axis shows chromosome 1H from short (left) to long (right) arm in base pair (bp). Red bars represent the candidate region harbouring the FCR resistant locus *Qcrs.cpi-1H*.

**Fig. 4.**
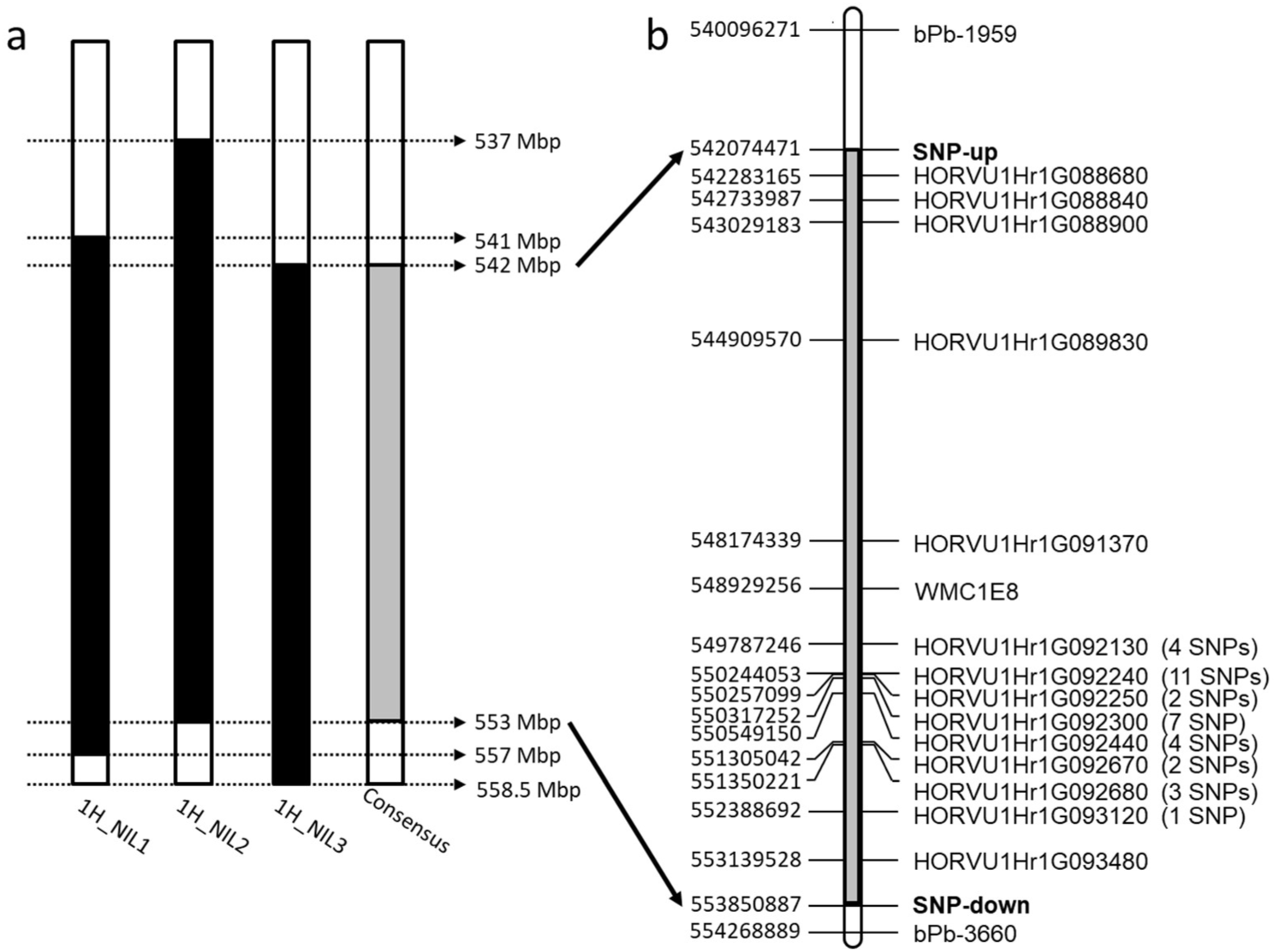
Physical distribution of DEGs within the consensus SNP-enriched region **a** Physical range of SNP-enriched regions. Black boxes indicate the regions defined by SNP within each 1H_NIL pair; the grey box represents for the consensus region. **b** Physical distribution of DEGs commonly detected from three comparisons within the consensus region. The initial QTL region was flanked by *bPb-1595* and *bPb-3660*. SNP-up/down indicate the borders of consensus region. The numbers of SNP identified within genes were bracketed

### DEGs with SNPs between the resistant and susceptible isolines targeting the *Qcrs.cpi-1H* locus

Based on the reference genome of barley cv. Morex, 266 high-confidence (HC) genes were identified within the common interval across three 1H_NIL pairs (Table S5). Among these HC genes, fifty-six carried SNPs and 14 were differentially expressed in one or more pairwise comparisons (Fig. 4b; Tables S2 and S3). Notably, five protein-coding genes were not only differentially expressed across the three NIL pairs but also carried SNPs led to changes in amino acids (Tables 3 and S3). These protein-coding genes should form the primary targets in identifying candidate genes underlying FCR resistance at this locus.

**Table 3.** Expression patterns of five DEGs bearing non-synonymous SNPs located in the interval harbouring the FCR resistant locus *Qcrs.cpi-1H*

| Gene ID | Gene Description <sup>a</sup> | Number of Non-synonymous SNPs | Expressed pattern |
| --- | --- | --- | --- |
| HORVU1Hr1G092130 | WRKYDNA-binding protein 23 | 1 | Upregulated in 3S isolines post Inoculation |
| HORVU1Hr1G092240 | Glucanendo-1,3-beta-glucosidase13 | 4 | Upregulated in 3R isolines post Inoculation |
| HORVU1Hr1G092250 | Receptor-like kinase | 1 | Upregulated in 3R and 3S isolines post Inoculation |
| HORVU1Hr1G092300 | Receptor-like kinase | 6 | Upregulated in 3R post Inoculation |
| HORVU1Hr1G092440 | P-loop containing nucleoside triphosphate hydrolases super family protein | 4 | Upregulated in 3S isolines post Inoculation |

<sup>a</sup>Gene descriptions were retrieved from the annotation file of the genome of barley cv. Morex

### Comparison of transcriptomic responses to *F. pseudograminearum* between 1H_NILs and 4H_NILs

To assess whether similarity in FCR resistance exists between 1H_NILs and 4H_NILs, we compared the transcriptomic profiles of *Qcrs.cpi-1H* obtained in this study with that for *Qcrs.cpi-4H* obtained from an earlier study [29]. As DEGs induced by *Fp*-inoculation for the latter were only identified from one of the 4H_NIL pairs (from comparisons R^M^_vs_R^S^ and S^M^_vs_S^S^), we compared its DEG datasets with those from each of the 1H_NIL pairs grouped as either ‘R’ (Fig. 5a) or ‘S’ (Fig. 5b) isolines. The correlations in DEG patterns between 1HL and 4HL datasets varied from 0.42 to 0.45 for the ‘R’ isolines and from 0.40 to 0.58 for the ‘S’ isolines. An overrepresentation analysis based on DEGs from the 1H_NILs detected a series of enriched GO terms strongly related to anti-oxidation (GO:0070279; GO:0030170; GO:0019842) and virulence detoxification (GO:0005506; GO:0020037; GO:0046906) pathways (Table S4). However, none of these enriched GO terms in 1H_NILs were detected in the 4H_NILs (not shown). We also compared the genome-wide distribution of DEGs mediated by *Qcrs.cpi-1H* with that from *Qcrs.cpi-4H* under *Fp*-inoculation (Fig. 6). The total number of DEGs in the ‘S’ isolines was higher than that in the ‘R’ isolines for NILs targeting both the 1HL and 4HL loci. However, the magnitudes of differential expression (i.e. value of fold-change) were higher for the DEGs from the 1H_NILs compared with those from the 4H_NILs.

**Fig. 5.**
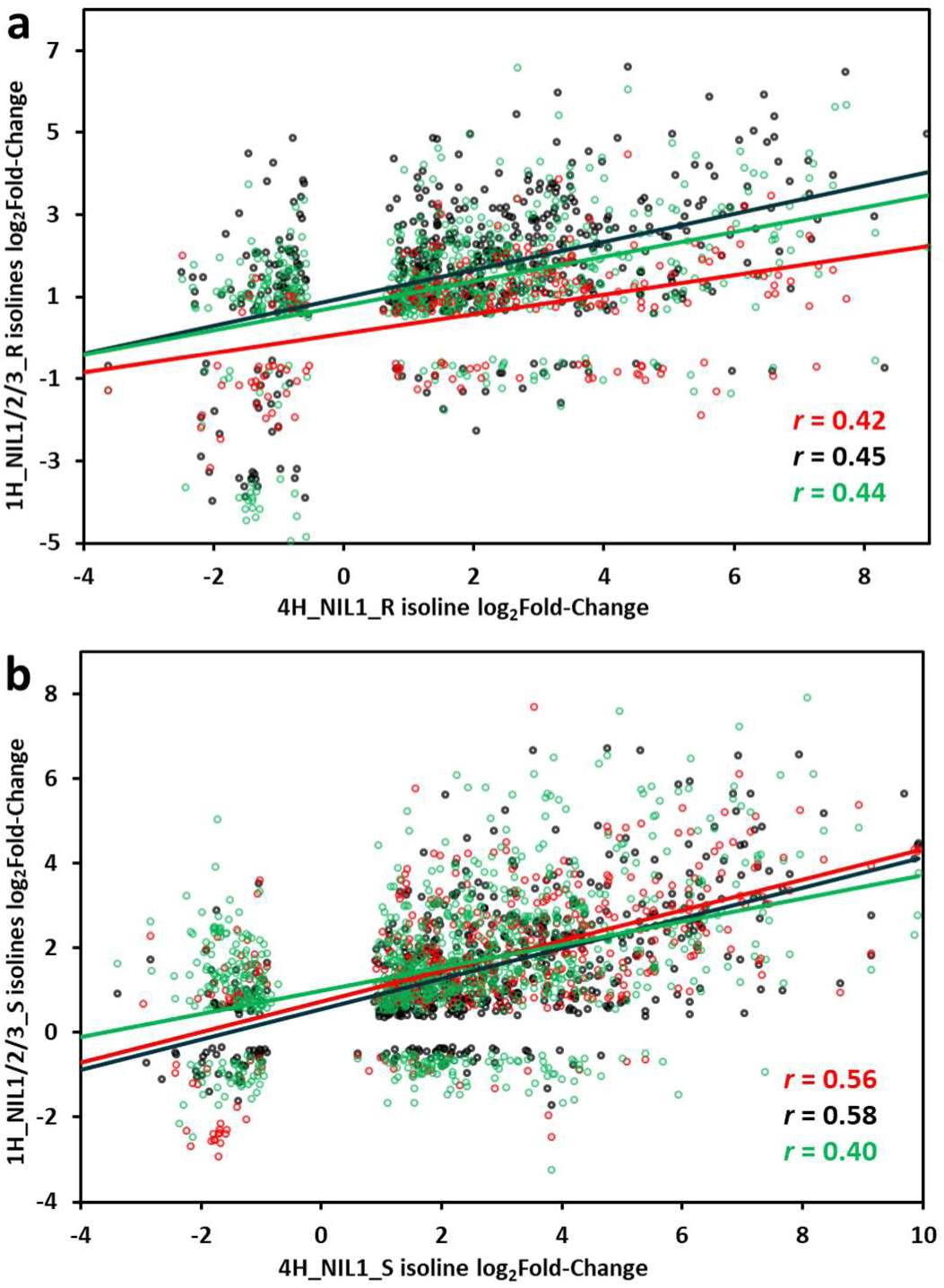
Comparison of log_2_fold-change values of *Fp*-induced DEGs between 4H_NILs (one pair) and 1H_NILs (three pairs), ‘**a**’ for difference between the ‘R’ lines and ‘**b’** for the ‘S’ lines. Results of the comparisons between the 4H-NIL pair with the three 1H-NIL pairs were differently coloured: 1H_NIL1 R/S in red, 1H_NIL2 R/S in black, and 1H_NIL3 R/S in green. Data points indicate comparison between a DEG in 4H_NIL1 R/S and its counterpart in 1H_NIL1 R/S (red), 1H_NIL2 R/S (black) and 1H_NIL3 R/S (green).

**Fig. 6.**
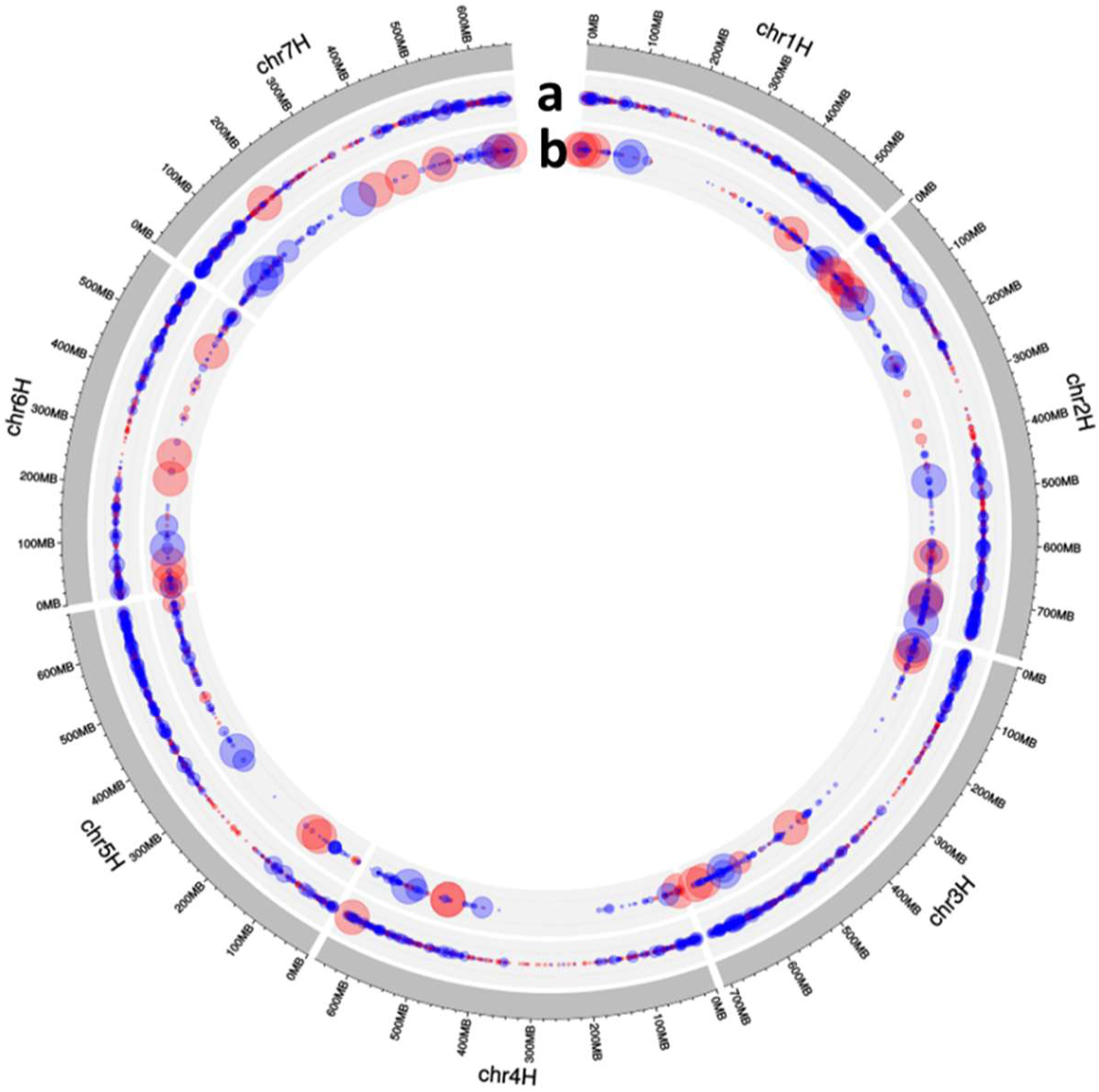
Genome-wide distributions of DEGs between ‘R’ and ‘S’ isolines under *Fp*-inoculation. The outmost circle represents the seven chromosomes (chr1H to chr7H) of the barley genome. DEGs identified in the comparison of R^I^_v_S^I^ from the three 4_NIL pairs (a) and those of the three pairs of 1H_NILs (b). Pink dots are for those genes with enhanced expression in the ‘R’ isolines, and blue dots are genes with enhanced expression in the ‘S’ isolines. Dot sizes represent absolute log_2_fold-changes of DEGs (the largest dot ≥ 11 log_2_FC).

## Discussion

FCR is a chronic disease for cereal production in semi-arid regions worldwide. It has long been recognised that breeding and growing resistant varieties have to form an integral part in the effect of effectively reducing damages from the disease. Previous studies also show that strong interactions between FCR severity and several characteristics including flowering time and plant height exist thus QTL detected from mapping populations need to be validated. In the study reported here, we successfully validated the QTL on chromosome arm 1HL by developing and assessing NILs targeting the locus. DEGs with SNPs shared by three pairs of the NILs further delineated the locus to an interval of about 11.0 Mbp. They would be invaluable for fine mapping the locus and cloning the gene(s) underlying its resistance. SNPs in several of the DEGs lead to amino acid changes and they would be primary targets in investigating the mechanism of FCR resistance.

It is of note that significant variation was found in the numbers of DEGs detected among the three pairs of NILs assessed. Previous studies showed that FCR development can be affected by various characteristics including plant height [11, 17, 21, 43] and flowering time [12, 16, 44]. Each of the NIL pairs used in this study was developed from a different heterozygous plant based on the profile of a single marker. This method ensured that different NIL pairs, including those from the same population, would have different genetic backgrounds. The different genetic backgrounds would lead to difference in FCR development at any given time point. In other words, although symptom of FCR infection was not visually observable for any of the NILs at 4 dpi when the samples for RNA-seq were taken, the advancement of FCR development among them must be different.

The interactions between FCR severity and other characteristics may also contributed to the difference in the effects of the 1HL locus between the use of NILs as described in this study and that based on QTL mapping [10]. In addition to the targeted trait, many other characteristics likely also segregate in populations routinely used for QTL mapping. They include populations of recombinant inbred lines and doubled haploid lines. In essence, a targeted locus is always assessed in different genetic backgrounds in QTL mapping studies, making its accurate assessment difficult. In the contrary, the two isolines forming each NIL pair differ mainly by the targeted locus. The fact that assessments for any characteristics can be carried out by comparing two isolines only must also contribute to the likelihood that more accurate assessment can be achieved by using NILs.

Of the DEGs with SNPs located in the interval harbouring the 1HL locus, several are known to be involved in plant-pathogen interaction. They include the two receptor-like kinase (RLK) genes which are involved in the immune systems in various plant species [45]. RLK locates on either the plasma or cytoplasmic membrane and are responsible for recognizing elicitor, usually small secreted protein, generated by pathogens. The perception of elicitor often triggers a fierce hypersensitive response (HR) which can cause programmed cell death [46]. Another one is the gene for glucanendo-1,3,-beta-glucosidase which plays an important role in defence against pathogen infection [47]. Its expression has been detected in the response to biotic stress in various plant species [48, 49]. The gene encoding a P-loop containing nucleoside triphosphate hydrolases (P-loop NTPase) protein is also among the DEGs with SNPs located in the targeted interval. Previous results showed that this gene negatively regulates plant defence response in both rice and *Arabidopsis* [50, 51]. Once bonded with ATP, *OsYchF1*, a P-loop NTPase in rice, contributes to resistance to biotic stress [52].

It is also of interesting to note that one of the DEGs with SNPs located in the targeted interval confers tolerance to drought. This is *HORVU1Hr1G092130* which codes a WRKY transcription factor which plays a key role in signalling in the defense response to biotic and abiotic stress [53, 54]. A homolog of *HORVU1Hr1G092130* in rice, *Os05g0583000* was strongly induced during drought response [55]. Over-expression of *Os05g0583000* coding sequence in *Arabidopsis* provided improved drought tolerance [56]. The presence of this gene related to drought tolerance is not a surprise as the relationship between drought stress and Fusarium crown rot severity in agricultural systems has been well documented. FCR causes severe yield loss mainly in semi-arid regions [1] and drought stress forms part of the procedures in FCR assay in both wheat [57, 58] and barley [10, 12, 13].

Comparison of transcriptomic results from NILs targeting *Qcrs.cpi-1H* and those targeting another FCR locus on 4HL showed that different mechanisms are likely involved in FCR resistance conferred by these two loci for three reasons: firstly, the correlation between DEG patterns from 1HL and 4HL studies were relatively low (on average 0.44 for ‘R’ group and 0.51 for ‘S’ group), especially considering that the correlation between *F. pseudograminearum*-induced transcriptomic profiles of *Brachypodium* and wheat sub-genomes could be 0.82-0.85 [59]. Secondly, the candidate genes obtained in this study have no functional overlap with genes in the fine-mapped interval of the 4HL FCR locus [30]. Thirdly, Habib et al. [29] reported that *Qcrs.cpi-4H* likely employed salicylic acid-mediated systemic defense signalling and triggered the synthesis of structural barriers to prevent pathogen infection. However, the results from this study indicated that the resistance regulated by *Qcrs.cpi-1H* likely involved anti-oxidation and DON detoxification pathways which also have been detected in the responses to *F. graminearum* in wheat [60, 61].

## Conclusions

In this study, we developed five pairs of NILs targeting the FCR resistance locus *Qcrs.cpi-1H*. Phenotyping these NIL found that the resistant allele at *Qcrs.cpi-1H* could significantly reduce FCR severity. Gene expression and SNP analysis of transcriptomic data derived from three pairs of the 1H_NILs delineated the *Qcrs.cpi-1H* locus into an about 11 Mbp interval containing 56 genes with SNP(s). Of these genes, five DEGs bearing non-synonymous SNPs form primary targets in identifying gene(s) underlying the *Qcrs.cpi-1H* locus. Lack of similarity between genes regulated between the 1HL and 4HL loci indicate the possible existence of different mechanisms in FCR resistance.

## Declarations

### Availability of supporting data and materials

The RNA sequences were available at the National Centre for Biotechnology Information (NCBI) with the accession number of PRJNA541021. The other supporting data were included as additional files.

### Author’s contribution

CL and MZ conceived and designed the experiments. SG and HA developed and assessed the NILs. SG, ZZ, JP conducted samples preparation and qRT-PCR validation. SG and JS performed the data analysis and SG drafted the manuscript. CL and other authors revised the manuscript. All authors read and approved the final manuscript.

## Acknowledgements

SG is grateful to University of Tasmania, Australia, and the China Scholarship Council for financial supports during the tenure of his PhD studentship. We are grateful to Drs Gao Lingling and Udaykumar Kage (both at CSIRO Agriculture and Food) for their constructive suggestions in preparing the manuscript.

## References

1. Chakraborty S, Liu C, Mitter V, Scott J, Akinsanmi O, Ali S, Dill-Macky R, Nicol J, Backhouse D, Simpfendorfer S: Pathogen population structure and epidemiology are keys to wheat crown rot and Fusarium head blight management. Australasian Plant Pathology 2006, 35(6):643–655.

2. Hogg A, Johnston R, Johnston J, Klouser L, Kephart K, Dyer A: Monitoring Fusarium crown rot populations in spring wheat residues using quantitative real-time polymerase chain reaction. Phytopathology 2010, 100(1):49–57.

3. Cook RJ: Fusarium foot rot of wheat and its control in the Pacific Northwest. Plant Dis 1980, 64(12):1061–1066.

4. Cook RJ: Management of wheat and barley root diseases in modern farming systems. Australasian Plant Pathology 2001, 30(2):119–126.

5. Kirkegaard J, Simpfendorfer S, Holland J, Bambach R, Moore K, Rebetzke G: Effect of previous crops on crown rot and yield of durum and bread wheat in northern NSW. Crop and Pasture Science 2004, 55(3):321–334.

6. Purss G: Studies of varietal resistance to crown rot of wheat caused by *Fusarium graminearum* Schw. Queensland Journal of Agricultural and Animal Sciences 1966, 23:475–498.

7. Zheng Z, Gao S, Zhou M, Yan G, Liu C: Enhancing Fusarium crown rot resistance by pyramiding large-effect QTL in common wheat (Triticum aestivum L.). Molecular Breeding 2017, 37(9):107.

8. Chen G, Habib A, Wei Y, Zheng Y-L, Shabala S, Zhou M, Liu C: Enhancing Fusarium crown rot resistance by pyramiding large-effect QTL in barley. Molecular breeding 2015, 35(1):26.

9. Liu C, Ogbonnaya FC: Resistance to Fusarium crown rot in wheat and barley: a review. Plant Breeding 2015, 134(4):365–372.

10. Chen G, Liu Y, Wei Y, McIntyre C, Zhou M, Zheng Y-L, Liu C: Major QTL for Fusarium crown rot resistance in a barley landrace. Theoretical and applied genetics 2013, 126(10):2511–2520.

11. Li HB, Zhou M, Liu CJ: A major QTL conferring crown rot resistance in barley and its association with plant height. Theoretical and Applied Genetics 2009, 118(5):903–910.

12. Chen G, Liu Y, Ma J, Zheng Z, Wei Y, McIntyre CL, Zheng Y-L, Liu C: A Novel and Major Quantitative Trait Locus for Fusarium Crown Rot Resistance in a Genotype of Wild Barley *(Hordeum spontaneum* L.). PloS one 2013, 8(3):e58040.

13. Gao S, Zheng Z, Hu H, Shi H, Ma J, Liu Y, Wei Y, Zheng Y, Zhou M, Liu C: A novel QTL conferring Fusarium crown rot resistance located on chromosome arm 6HL in barley. BioRxiv 2019:537605.

14. Zheng Z, Ma J, Stiller J, Zhao Q, Feng Q, Choulet F, Feuillet C, Zheng Y-L, Wei Y, Han B: Fine mapping of a large-effect QTL conferring Fusarium crown rot resistance on the long arm of chromosome 3B in hexaploid wheat. BMC genomics 2015, 16(1):850.

15. Yan W, Li H, Cai S, Ma H, Rebetzke G, Liu C: Effects of plant height on type I and type II resistance to fusarium head blight in wheat. Plant Pathology 2011, 60(3):506–512.

16. Liu Y, Zheng YL, Wei Y, Zhou M, Liu C: Genotypic differences to crown rot caused by Fusarium pseudograminearum in barley (Hordeum vulgare L.). Plant Breeding 2012, 131(6):728–732.

17. Liu YX, Yang XM, Ma J, Wei YM, Zheng YL, Ma HX, Yao JB, Yan GJ, Wang YG, Manners JM: Plant height affects Fusarium crown rot severity in wheat. Phytopathology 2010, 100(12):1276–1281.

18. Chen G, Li H, Zheng Z, Wei Y, Zheng Y, McIntyre C, Zhou M, Liu C: Characterization of a QTL affecting spike morphology on the long arm of chromosome 3H in barley (Hordeum vulgare L.) based on near isogenic lines and a NIL-derived population. Theoretical and applied genetics 2012, 125(7):1385–1392.

19. Liu X, Liu C: Effects of drought-stress on Fusarium crown rot development in Barley. PloS one 2016, 11(12):e0167304.

20. Pumphrey MO, Bernardo R, Anderson JA: Validating the QTL for Fusarium head blight resistance in near-isogenic wheat lines developed from breeding populations. Crop Science 2007, 47(1):200–206.

21. Chen G, Yan W, Liu Y, Wei Y, Zhou M, Zheng Y-L, Manners JM, Liu C: The non-gibberellic acid-responsive semi-dwarfing gene *uzu* affects Fusarium crown rot resistance in barley. BMC plant biology 2014, 14(1):22.

22. Ma J, Yan GJ, Liu CJ: Development of near-isogenic lines for a major QTL on 3BL conferring Fusarium crown rot resistance in hexaploid wheat. Euphytica 2012, 183(2):147–152.

23. Habib A, Shabala S, Shabala L, Zhou M, Liu C: Near-isogenic lines developed for a major QTL on chromosome arm 4HL conferring Fusarium crown rot resistance in barley. Euphytica 2016, 209(3):555–563.

24. Mortazavi A, Williams BA, McCue K, Schaeffer L, Wold B: Mapping and quantifying mammalian transcriptomes by RNA-Seq. Nature methods 2008, 5(7):621.

25. Wang Z, Gerstein M, Snyder M: RNA-Seq: a revolutionary tool for transcriptomics. Nature reviews genetics 2009, 10(1):57.

26. Blencowe BJ, Ahmad S, Lee LJ: Current-generation high-throughput sequencing: deepening insights into mammalian transcriptomes. Genes & development 2009, 23(12):1379–1386.

27. Cavanagh CR, Chao S, Wang S, Huang BE, Stephen S, Kiani S, Forrest K, Saintenac C, Brown-Guedira GL, Akhunova A: Genome-wide comparative diversity uncovers multiple targets of selection for improvement in hexaploid wheat landraces and cultivars. Proceedings of the national academy of sciences 2013, 110(20):8057–8062.

28. Ma J, Stiller J, Zhao Q, Feng Q, Cavanagh C, Wang P, Gardiner D, Choulet F, Feuillet C, Zheng Y-L: Transcriptome and Allele Specificity Associated with a 3BL Locus for Fusarium Crown Rot Resistance in Bread Wheat. 2014.

29. Habib A, Powell JJ, Stiller J, Liu M, Shabala S, Zhou M, Gardiner DM, Liu C: A multiple near isogenic line (multi-NIL) RNA-seq approach to identify candidate genes underpinning QTL. Theoretical and applied genetics 2018, 131(3):613–624.

30. Jiang Y, Habib A, Zheng Z, Zhou M, Wei Y, Zheng Y-L, Liu C: Development of tightly linked markers and identification of candidate genes for Fusarium crown rot resistance in barley by exploiting a near-isogenic line-derived population. Theoretical and Applied Genetics 2019, 132(1):217–225.

31. Tuinstra M, Ejeta G, Goldsbrough P: Heterogeneous inbred family (HIF) analysis: a method for developing near-isogenic lines that differ at quantitative trait loci. Theoretical and Applied Genetics 1997, 95(5-6):1005–1011.

32. Zheng Z, Wang H, Chen G, Yan G, Liu C: A procedure allowing up to eight generations of wheat and nine generations of barley per annum. Euphytica 2013, 191(2):311–316.

33. Akinsanmi OA, Mitter V, Simpfendorfer S, Backhouse D, Chakraborty S: Identity and pathogenicity of *Fusarium* spp. isolated from wheat fields in Queensland and northern New South Wales. Crop and Pasture Science 2004, 55(1):97–107.

34. Li X, Liu C, Chakraborty S, Manners JM, Kazan K: A simple method for the assessment of crown rot disease severity in wheat seedlings inoculated with *Fusarium pseudograminearum*. Journal of Phytopathology 2008, 156(11-12):751–754.

35. Trapnell C, Roberts A, Goff L, Pertea G, Kim D, Kelley DR, Pimentel H, Salzberg SL, Rinn JL, Pachter L: Differential gene and transcript expression analysis of RNA-seq experiments with TopHat and Cufflinks. Nature protocols 2012, 7(3):562.

36. Mascher M, Gundlach H, Himmelbach A, Beier S, Twardziok SO, Wicker T, Radchuk V, Dockter C, Hedley PE, Russell J: A chromosome conformation capture ordered sequence of the barley genome. Nature 2017, 544(7651):427.

37. Yu Y, Ouyang Y, Yao W: shinyCircos: an R/Shiny application for interactive creation of Circos plot. Bioinformatics 2017, 34(7):1229–1231.

38. Oliveros J: An interactive tool for comparing lists with Venn’s diagrams (2007–2015). In.; 2018.

39. Stephen S, Cullerne D, Spriggs A, Helliwell C, Lovell D, Taylor J: BioKanga: a suite of high performance bioinformatics applications. In.: preparation; 2012.

40. Cingolani P, Platts A, Wang LL, Coon M, Nguyen T, Wang L, Land SJ, Lu X, Ruden DM: A program for annotating and predicting the effects of single nucleotide polymorphisms, SnpEff: SNPs in the genome of Drosophila melanogaster strain w1118; iso-2; iso-3. Fly 2012, 6(2):80–92.

41. Conesa A, Götz S, García-Gómez JM, Terol J, Talón M, Robles M: Blast2GO: a universal tool for annotation, visualization and analysis in functional genomics research. Bioinformatics 2005, 21(18):3674–3676.

42. Du Z, Zhou X, Ling Y, Zhang Z, Su Z: agriGO: a GO analysis toolkit for the agricultural community. Nucleic acids research 2010, 38(Suppl_2):W64–W70.

43. Bai Z, Liu C: Histological Evidence for Different Spread of F usarium crown rot in Barley Genotypes with Different Heights. Journal of Phytopathology 2015, 163(2):91–97.

44. Liu Y, Ma J, Yan W, Yan G, Zhou M, Wei Y, Zheng Y, Liu C: Different tolerance in bread wheat, durum wheat and barley to Fusarium crown rot disease caused by Fusarium pseudograminearum. Journal of Phytopathology 2012, 160(7-8):412–417.

45. Marone D, Russo M, Laidò G, De Leonardis A, Mastrangelo A: Plant nucleotide binding site– leucine-rich repeat (NBS-LRR) genes: active guardians in host defense responses. International journal of molecular sciences 2013, 14(4):7302–7326.

46. Krattinger SG, Keller B: Molecular genetics and evolution of disease resistance in cereals. New Phytologist 2016, 212(2):320–332.

47. Beffa RS, Neuhaus J-M, Meins F: Physiological compensation in antisense transformants: specific induction of an” ersatz” glucan endo-1, 3-beta-glucosidase in plants infected with necrotizing viruses. Proceedings of the National Academy of Sciences 1993, 90(19):8792–8796.

48. Faghani E, Gharechahi J, Komatsu S, Mirzaei M, Khavarinejad RA, Najafi F, Farsad LK, Salekdeh GH: Comparative physiology and proteomic analysis of two wheat genotypes contrasting in drought tolerance. Journal of proteomics 2015, 114:1–15.

49. Su Y, Wang Z, Liu F, Li Z, Peng Q, Guo J, Xu L, Que Y: Isolation and characterization of ScGluD2, a new sugarcane beta-1, 3-glucanase D family gene induced by Sporisorium scitamineum, ABA, H2O2, NaCl, and CdCl2 stresses. Frontiers in plant science 2016, 7:1348.

50. Cheung MY, Li MW, Yung YL, Wen CQ, Lam HM: The unconventional P-loop NTPase OsYchF1 and its regulator OsGAP1 play opposite roles in salinity stress tolerance. Plant, cell & environment 2013, 36(11):2008–2020.

51. Cheung M-Y, Xue Y, Zhou L, Li M-W, Sun SS-M, Lam H-M: An ancient P-loop GTPase in rice is regulated by a higher plant-specific regulatory protein. Journal of Biological Chemistry 2010, 285(48):37359–37369.

52. Cheung M-Y, Li X, Miao R, Fong Y-H, Li K-P, Yung Y-L, Yu M-H, Wong K-B, Chen Z, Lam H-M: ATP binding by the P-loop NTPase OsYchF1 (an unconventional G protein) contributes to biotic but not abiotic stress responses. Proceedings of the National Academy of Sciences 2016, 113(10):2648–2653.

53. Eulgem T, Rushton PJ, Robatzek S, Somssich IE: The WRKY superfamily of plant transcription factors. Trends in plant science 2000, 5(5):199–206.

54. Birkenbihl RP, Kracher B, Roccaro M, Somssich IE: Induced genome-wide binding of three Arabidopsis WRKY transcription factors during early MAMP-triggered immunity. The Plant Cell 2017, 29(1):20–38.

55. Shin S-J, Ahn H, Jung I, Rhee S, Kim S, Kwon H-B: Novel drought-responsive regulatory coding and non-coding transcripts from Oryza Sativa L. Genes & Genomics 2016, 38(10):949–960.

56. Song Y, Jing S, Yu D: Overexpression of the stress-induced OsWRKY08 improves osmotic stress tolerance in Arabidopsis. Chinese Science Bulletin 2009, 54(24):4671–4678.

57. Ma J, Li HB, Zhang CY, Yang XM, Liu YX, Yan GJ, Liu CJ: Identification and validation of a major QTL conferring crown rot resistance in hexaploid wheat. Theoretical and Applied Genetics 2010, 120(6):1119–1128.

58. Zheng Z, Kilian A, Yan G, Liu C: QTL Conferring Fusarium Crown Rot Resistance in the Elite Bread Wheat Variety EGA Wylie. PloS One 2014, 9(4):e96011.

59. Powell JJ, Carere J, Sablok G, Fitzgerald TL, Stiller J, Colgrave ML, Gardiner DM, Manners JM, Vogel JP, Henry RJ: Transcriptome analysis of Brachypodium during fungal pathogen infection reveals both shared and distinct defense responses with wheat. Scientific reports 2017, 7(1):17212.

60. Lemmens M, Scholz U, Berthiller F, Dall’Asta C, Koutnik A, Schuhmacher R, Adam G, Buerstmayr H, Mesterházy Á, Krska R: The ability to detoxify the mycotoxin deoxynivalenol colocalizes with a major quantitative trait locus for Fusarium head blight resistance in wheat. Molecular plant-microbe interactions 2005, 18(12):1318–1324.

61. Li X, Shin S, Heinen S, Dill-Macky R, Berthiller F, Nersesian N, Clemente T, McCormick S, Muehlbauer GJ: Transgenic wheat expressing a barley UDP-glucosyltransferase detoxifies deoxynivalenol and provides high levels of resistance to Fusarium graminearum. Molecular Plant-Microbe Interactions 2015, 28(11):1237–1246.

