## Supplementary material for "Validation and delineation of a locus conferring Fusarium crown rot resistance on 1HL in barley by analysing transcriptomes from multiple pairs of near isogenic lines": Figure S1

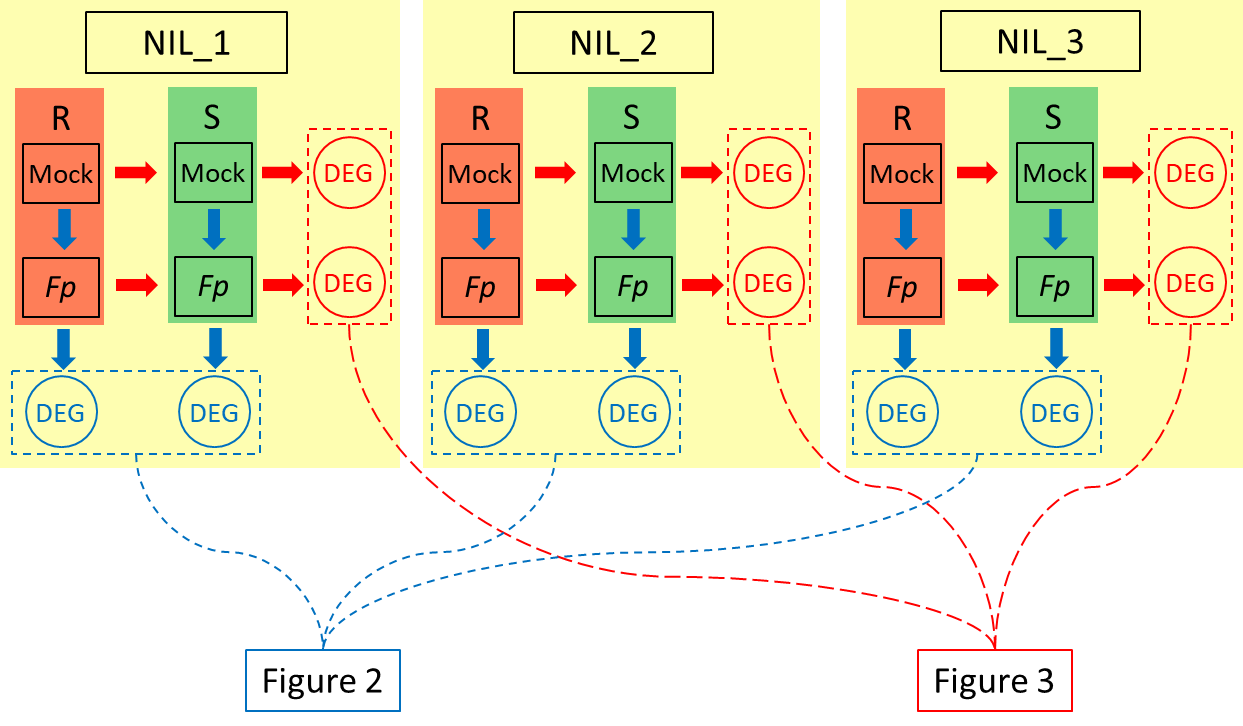

**Supplementary Figure 1.** Schematic showing the experimental design for differential gene expression analysis.
